# p53 induces senescence in the unstable progeny of aneuploid cells

**DOI:** 10.1101/818112

**Authors:** Maybelline Giam, Cheng Kit Wong, Jun Siong Low, Matteo Sinelli, Oliver Dreesen, Giulia Rancati

## Abstract

Aneuploidy is the condition of having an imbalanced karyotype, which is strongly associated with tumor initiation, evolution, and acquisition of drug-resistant features, possibly by generating heterogeneous populations of cells with distinct genotypes and phenotypes. Multicellular eukaryotes have therefore evolved a range of extrinsic and cell-autonomous mechanisms for restraining proliferation of aneuploid cells, including activation of the tumor suppressor protein p53. However, accumulating evidence indicates that a subset of aneuploid cells can escape p53-mediated growth restriction and continue proliferating *in vitro*. Here we show that such aneuploid cell lines display a robust modal karyotype and low frequency of chromosomal aberrations despite ongoing chromosome instability. Indeed, while these aneuploid cells are able to survive for extended periods *in vitro*, their chromosomally unstable progeny remain subject to p53-induced senescence and growth restriction, leading to subsequent elimination from the aneuploid pool. This mechanism helps maintain low levels of heterogeneity in aneuploid populations and may prevent detrimental evolutionary processes such as cancer progression and development of drug resistance.

## Introduction

Aneuploidy profoundly alters cell phenotype and is associated with developmental abnormalities and growth retardation at the whole organism level [1]. Despite impairing the proliferative capacity of individual cells, aneuploidy correlates with cancer evolution and therapy resistance [2], possibly due to the fitness advantages this confers under specific environmental and genetic conditions [3-7]. Moreover, since aneuploidy has been shown to cause genetic and chromosomal instability (CIN) in both yeast and mammalian cells [8-11], this state may promote the generation of heterogeneous populations that support cancer development and tumor acquisition of drug resistant traits [7,12,13]. Given this risk of potential pathological changes, multicellular organisms have evolved a range of different strategies to curb the proliferation of aneuploid cells. One such cell-autonomous strategy is centered on the transcription factor and tumor suppressor protein p53, which relocates to the cell nucleus to promote specific transcriptional responses to a variety of stress stimuli [14]. Previous reports have suggested that activation of p53 can be triggered directly via the stress kinase p38 [15] or mediated by DNA damage or accumulation of reactive oxygen species (ROS) following chromosome mis-segregation [16-18]. Other data suggest that phosphorylation of histone H3 on mis-segregated chromosomes is sufficient to stabilize p53 independently of key DNA-damaging signaling proteins [19]. However, accumulating evidence indicates that aneuploid cells can escape p53 signaling, and aneuploid cell lines can be established *in vitro* [11,18,20]. These aberrant aneuploid cells produce pro-inflammatory signals and can thus be actively removed by the host immune system *in vivo* [20]. While, previous reports have focused on mechanisms of growth inhibition following the first few cell divisions after chromosome mis-segregation in euploid populations, here we investigated whether cell-intrinsic mechanisms can prevent proliferation of the aberrant progeny of established aneuploid lines. We now report that cell-autonomous mechanisms of growth restriction are indeed capable of inducing senescence and halting cell cycle progression of karyotypically unstable daughters of aneuploid cells. These observations suggest that distinct control mechanisms limit the proliferation of aneuploid cells and the generation of cell-to-cell heterogeneity at different stages and have direct relevance for efforts to understand tumor cell evolution and improve treatment strategies in the cancer clinic.

## Results and Discussion

### Human cell lines can escape autonomous growth inhibition and become aneuploid *in vitro*

We sought to test whether cell-autonomous mechanisms of growth restriction can impair proliferation in the unstable progeny of aneuploid cells. To first generate aneuploid populations, we performed single-cell clonal amplification of the non-transformed retinal pigment epithelial RPE-1 cell line. The parental RPE-1 cells displayed marked genome stability and a robust modal diploid karyotype (Fig 1A, RPE + DMSO), while spectral karyotyping (SKY) confirmed that genomic aberrations were absent (excepting a known X-chromosome derivative [t(X;10)], Fig EV1A-D). To generate random whole-chromosome aneuploidies, synchronized metaphase-enriched RPE-1 cells were briefly exposed to reversine (Fig EV1E; see Materials and Methods for experimental details), which inhibits the spindle assembly checkpoint (SAC) protein kinase Mps1 [21]. Inhibition of Mps1 licenses metaphase to anaphase transition even in presence of chromosome attachment or positioning defects [22], thus generating daughter cells with random aneuploidies. Since Mps1 inhibition does not delay cell cycle progression, reversine treatment avoids the p53-dependent G1 arrest typically observed after prolonged mitotic delay (as known to be associated with other aneuploidy induction methods; [23]). To quantify the experimental induction of aneuploidy, chromosome counting was performed on metaphase spreads harvested 20 hours after initial reversine treatment. This approach allows sufficient time for cells that divided during the reversine exposure to restart cell cycle progression and reach metaphase by the time of harvesting (Fig EV1E). Following reversine treatment, more than half of the RPE cells had either gained or lost chromosomes relative to control cells treated with vehicle only (Fig 1A). SKY analysis confirmed the induction of random whole chromosome gains and losses with no evidence of translocation events (Fig 1B), suggesting that short exposure to reversine does not cause DNA damage or structural aberrations. Consistent with these findings, there was no significant increase in co-localization of γH2Ax and 53BP1 DNA repair foci following reversine treatment (Fig EV1F). Despite robust induction of aneuploidy by reversine treatment (47.4%, RPE+REV, Fig 1A), after single-cell cloning we were only able to find three aneuploid RPE-1 cell lines from a total of 111 viable clones analyzed (2.7%, Fig 1C), suggesting that only a minority of aneuploid cells can escape cell-autonomous growth restriction. We next performed G-banding analyses on all 3 aneuploid cell lines and 1 diploid control clone (RPE^46^) retrieved after single-cell amplification of reversine-treated RPE cells. Chromosome counting showed that both the aneuploid and diploid lines exhibited a robust modal karyotype with low degree of numerical and structural instability (Fig 1D). The karyotypes of the aneuploid cell lines with modal chromosome number 47, 48 and 49 were determined to be trisomy 12 (RPE^+12^), tetrasomy 18 (RPE^+18+18^) and trisomy 2, 12 and 19 (RPE^+2+12+19^), respectively, with no sign of chromosomal translocation (Fig 1E). Moreover, the karyotype of the RPE^46^ cell line was confirmed to be diploid with no signs of structural aberrations (Fig 1D and E). To verify whether the aneuploid cell lines retained a functional p53 pathway, these were subsequently exposed to the DNA-damaging agent doxorubicin. As shown in Figure EV1G, both p53 and downstream target p21 were stabilized in response to doxorubicin exposure, suggesting that these aneuploid cell lines were capable of proliferating despite the presence of an intact p53 signaling cascade.

**Figure 1.**
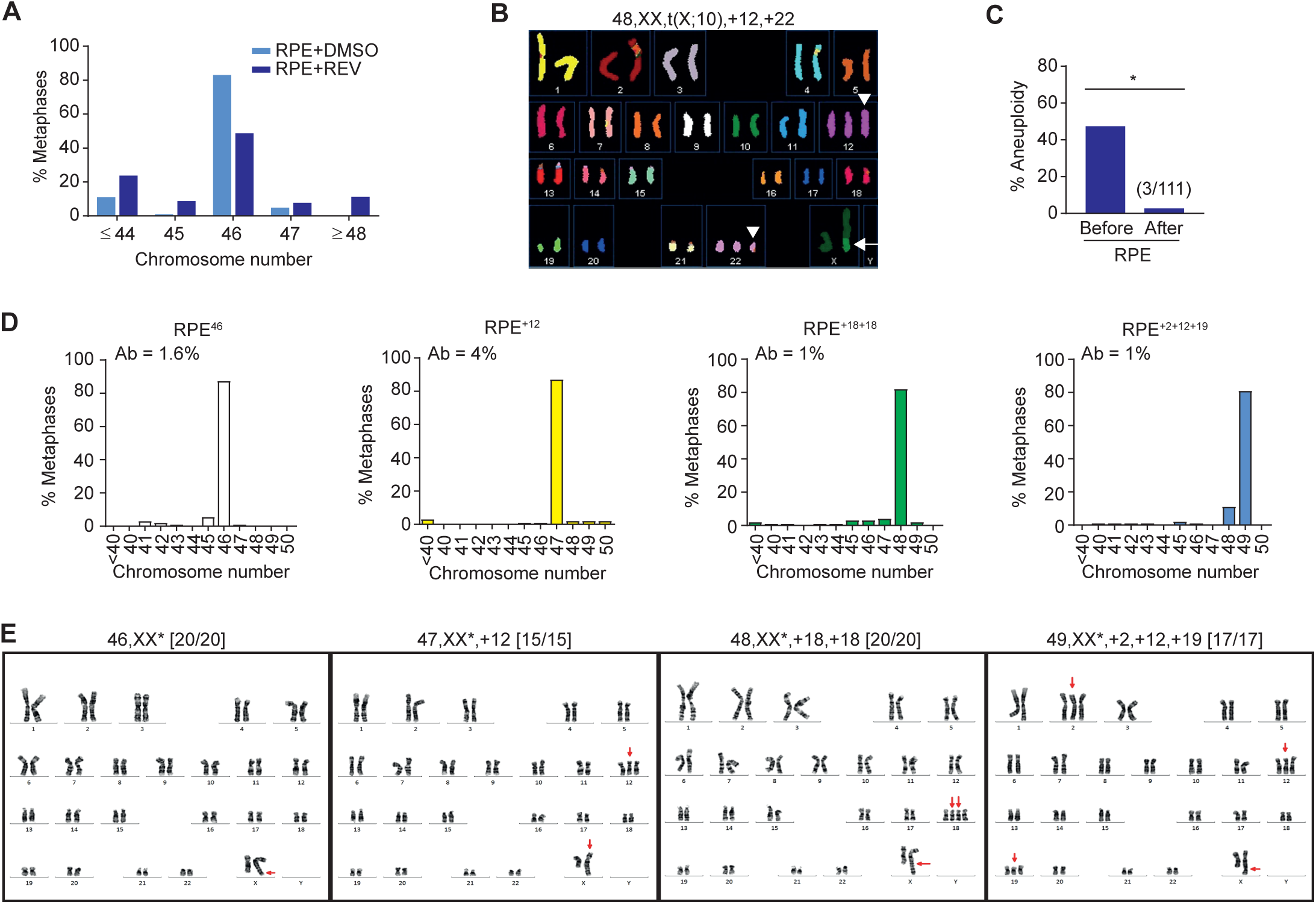
Aneuploid cells can survive *in vitro*. A Chromosome counts from metaphase spreads of RPE cells 20h after treatment with reversine (REV) or DMSO vehicle-only control (n=50; shown is the average of two independent replicates). B Representative spectral karyotype of an aneuploid RPE cell after reversine treatment. White arrow: t(X:10) present in the background (Fig EV1A-D). White arrowheads: chromosome gains after reversine treatment. C Percentage of aneuploid cells detected before and after single-cell clonal amplification; clones were classified aneuploid if their modal chromosome number calculated from 20 metaphase spreads ≠ 46 (*p< 0.05 by Fisher’s exact test). Polyploid cells with chromosome number >65 were seldom found D,E Chromosome counts (D) and G-banding karyotypes (E) of a diploid (RPE^46^) and three aneuploid lines recovered after single-cell clonal amplification of reversine-treated RPE-1 cells. (D) Histograms represent the distribution of chromosome number per cell from metaphase spreads. Percentage of cells with structurally aberrant chromosomes (Ab) is shown top left; n=50 metaphase spreads; shown is the average of two independent replicates. (E) Gained chromosomes and the t(X:10) translocation are indicated with red arrows; karyotypes (with derivative X chromosomes labelled with asterisks) are indicated at the top of the panels; karyotypic frequency is indicated in the brackets.

### Aneuploid cell lines display stable karyotypes despite chromosomal instability

Aneuploid karyotypes display different degrees of genome instability between model systems [8-11]. We therefore proceeded to test whether karyotype stability of the aneuploid lines obtained here was due to a high overall level of chromosomal integrity or could be attributed to growth restriction of the aberrant daughter cells. To this end, we used time-lapse imaging to visualize chromosome segregation during mitosis in both the diploid and aneuploid RPE cell lines (Fig 2A and B). As expected, the diploid RPE^46^ clone showed low rates of chromosome mis-segregation (Fig 2B). While chromosome mis-segregation was not significantly altered in RPE^+12^ cells, the RPE^+18+18^ and RPE^+2+12+19^ lines displayed increased occurrence of mitotic defects, including lagging chromosomes and bridges during anaphase (Fig 2B and Movie EV1). These defects did not confer any lengthening of mitosis, but correlated with an increase in the fraction of cells exhibiting micronuclei (MN) (Fig 2C and D). Micronuclei are extra-nuclear bodies that are typically formed as a consequence of chromosome mis-segregation / lagging / bridges. Therefore their presence likely identifies the products of aberrant mitosis (although a small fraction of MN-cells might also have undergone improper chromosome segregation without formation of micronuclei). To independently confirm the observed chromosome instability, RPE^+18+18^ aneuploid cells were treated with dihydrocytochalasin B (DCB) to disrupt cytokinesis followed by FISH labelling using specific probes to identify chromosomes 13, 18 and 21 in the binucleated cells (Fig 2E and F). This method reveals the reciprocal distribution of labeled chromosomes between daughter nuclei immediately after chromosome segregation and can be applied to the analysis of several hundred cells in tandem [10]. As shown in Fig 2E and F, RPE^+18+18^ cells displayed a significant increase in chromosome mis-segregation rates, consistent with the results of our live-cell imaging analysis. Moreover, aneuploid cell lines displayed a significant increase in DNA damage as indicated by co-localization of γH2Ax and 53BP1 foci per cell (Fig 2G and H).

**Figure 2.**
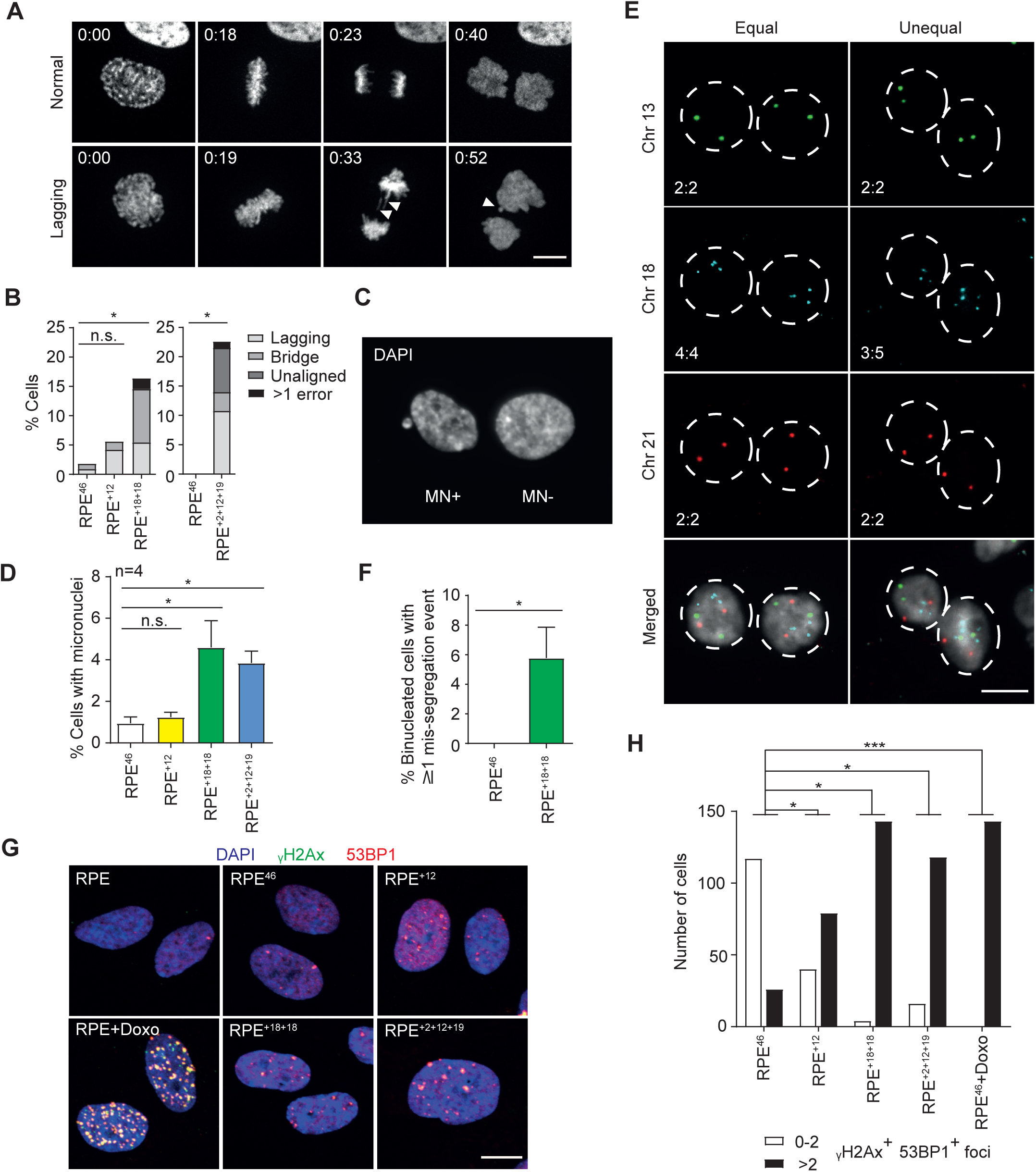
Aneuploid RPE cell lines display ongoing genome instability despite a stable modal karyotype. A Time-lapse microscopy montage of H2B-mCherry RPE^+18+18^ cells undergoing mitosis. Top: normal mitosis; bottom: aberrant mitosis (presence of lagging chromosomes and micronuclei, white arrowheads). Time frames are indicated as hr:min after Nuclear Envelope Breakdown (NEB, t = 0). Representative movies are also available as supplementary (Movie EV1). B Quantification of mitotic errors in the indicated H2B-mCherry RPE clones from the time-lapse movies. RPE^+2+12+19^ and another set of control cells were analyzed on IX83 Olympus (n=86-167; *p< 0.05 by Fisher’s exact test). C,D Representative image of cells containing (MN+) or lacking (MN-) micronuclei (C) and percentage of cells containing at least one micronucleus in the indicated lines (D). Shown are the average and SEM of four independent experiments (n≥1000; *p<0.05 by Student’s t test). E Representative images of chromosome-specific FISH hybridization in RPE^+18+18^ cell line after DCB-induced cytokinesis failure. Equal or unequal segregation: left and right columns, respectively. Chromosome numbers in the two daughter nuclei are shown bottom left. F Quantification of cells displaying at least one mis-segregation event as identified in (E). Data are presented as mean ± S.E.M. of three independent experiments (n=500; *p<0.05 by Student’s t test). G Representative images of immunolabeling with γH2Ax and 53BP1 in the indicated cell lines and conditions (Doxo = doxorubicin 400nM, 4 h); γH2Ax, 53BP1 and DAPI signals are overlaid. H Quantification of γH2Ax and 53BP1 positive foci in the indicated lines and conditions (n>100 cells). The experiment was repeated twice with qualitatively similar results (*p< 0.05 and ***p< 0.001 by Fisher’s exact test).

To verify our findings, we next analyzed the karyotypes and chromosome mis-segregation rates of primary fibroblast lines derived from one diploid (FIB^46^) and three trisomic human patients (trisomic 13: FIB^+13^, trisomic 18: FIB^+18^ and trisomic 21: FIB^+21^). We performed chromosome counting on all lines as well as multi-colour FISH (MFISH) karyotypic analysis of FIB^+13^ cells, which confirmed that all three aneuploid fibroblast lines displayed robust modal karyotypes with no sign of structural aberrations (Fig 3A-E). Quantification of chromosome mis-segregation after DCB-induced cytokinesis failure also revealed a significant increase in chromosome instability in FIB^+13^ and FIB^+21^ lines (Fig 3F and G). However, no significant increase in DNA damage was observed under these conditions, suggesting that there is no ongoing nucleotide instability in these lines. Stabilization of p53 and p21 in response to doxorubicin treatment was also consistently observed in the human patient-derived fibroblast lines (Fig EV1H), suggesting that these aneuploid populations featured intact p53 signaling.

**Figure 3.**
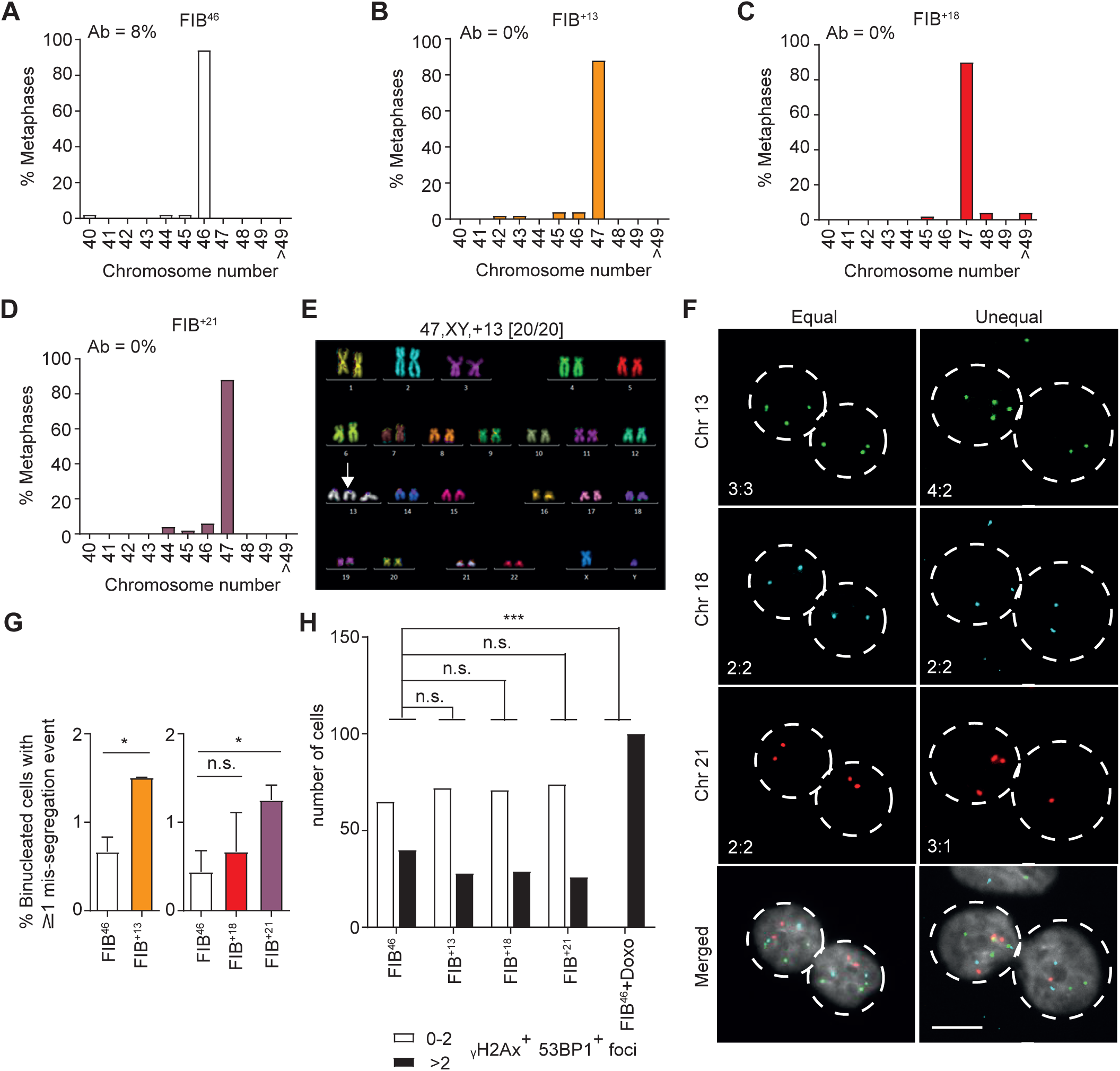
Primary aneuploid fibroblast cell lines are chromosomally unstable but maintain a stable karyotype. A-D Chromosome counting from metaphase spreads of the indicated euploid and aneuploid primary fibroblasts (n=50); percentages of cells with structurally aberrant chromosomes (Ab) are shown top left. E Representative pseudo-coloured image from M-FISH hybridization karyotyping of a FIB^+13^ cell. Karyotype is indicated on top with the number of cells analyzed in brackets. White arrow: trisomic chromosome. F Representative images of chromosome-specific FISH hybridization in FIB^+13^ cells after DCB-induced cytokinesis failure. Equal or unequal segregation: left and right column, respectively. Chromosome numbers in the two daughter nuclei are shown bottom left (scale bar: 10µm). G Percentage of cells displaying at least one mis-segregation event in the indicated cell lines. Data are presented as mean ± S.E.M. of three independent experiments (n≥522); *p<0.05 by Student’s t test. H Quantification of γH2Ax and 53BP1 positive foci in the indicated lines and conditions (Doxo = doxorubicin 400nM, 4 h) after immunoblotting staining (n>100 cells). The experiment was repeated twice with qualitatively similar results (***p< 0.001 by Fisher’s exact test).

Taken together, these results showed that aneuploid RPE cells and primary human fibroblasts display increased chromosome instability and ongoing DNA damage despite their robust modal karyotypes and lack of structural aberrations. These observations suggest that cell-autonomous mechanisms likely restrain the proliferation of aberrant daughter cells even after failing to limit growth of the aneuploid mother.

### Cell cycle arrest and senescence following chromosome mis-segregation

While the majority of the aneuploid cell lines under study displayed increased rates of chromosome segregation errors and DNA damage, we did not observe any increase in karyotypic heterogeneity and/or structural aberrations when analyzing metaphase spreads. A possible explanation for this observation could be that progeny arising from improper division of aneuploid cells are arrested in G1, fail to progress to mitosis, and therefore cannot be detected by metaphase spread analysis. To test this hypothesis, we compared the level of karyotypic heterogeneity in interphase versus metaphase for both aneuploid and diploid control RPE cell lines using FISH staining with locus-specific probes for chromosomes 13 and 18 (Fig 4A and B). We observed that copy number alterations in chromosomes 13 and 18 were significantly more common among interphase relative to metaphase cells in RPE^+18+18^ and RPE^+2+12+19^ aneuploid lines (Fig 4A and B), consistent with their ongoing chromosome mis-segregation (Fig 2A-F). This finding supports our hypothesis that cell-autonomous mechanisms restrain proliferation of the aberrant progeny of aneuploid cells during interphase.

**Figure 4.**
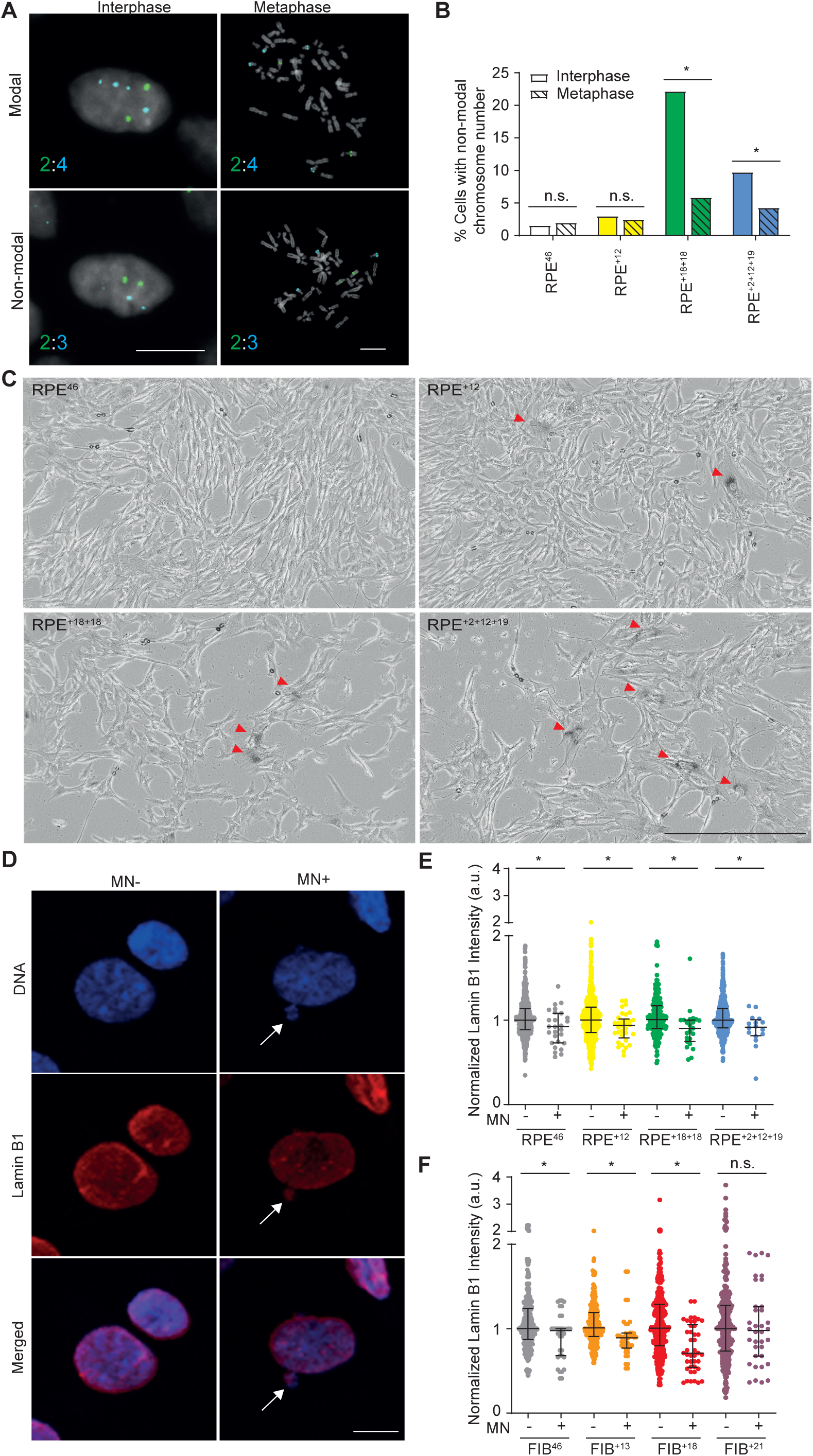
Cell cycle arrest following chromosome mis-segregation. A Representative images of interphase or metaphase FISH using specific probes for chromosomes 13 (green, 2 copies) and 18 (aqua, 4 copies) in the RPE^+18+18^ cell line. Chromosome counts per cell are shown bottom left. Top and bottom row: cells displaying modal or non-modal distribution of chromosomes 13 and 18, respectively (scale bar: 10µm). B Quantification of interphase (solid) or metaphase (pattern) cells with non-modal distribution of chromosomes 13 and 18 in the indicated cell lines. A total of 300-450 interphase and 85-200 metaphase cells were scored (*p<0.05 by Fisher’s exact test). C Representative images of β-galactosidase assay performed on the indicated cell lines (scale bar: 400µm). Red arrowhead indicate senescence cells. D Representative images of Lamin B1 immunofluorescence in RPE^+18+18^ cells with (MN+) or without (MN-) micronuclei (indicated by the white arrow). DNA was stained with DAPI (scale bar: 10µm). E,F Quantification of normalized Lamin B1 intensity in MN- and MN+ cells from the indicated cell lines. A total of 16-42 MN+ and 235-756 MN-cells were quantified and two biological replicates were performed with qualitatively similar results. Error bars represent the median and interquartile ranges (*p<0.05 by Mann-Whitney test).

Given that we did not observe cell death by apoptosis (Fig EV2), we next tested if the daughter cells generated by aberrant mitosis instead become permanently arrested in G1. Consistently, we found that senescence-associated β-galactosidase (SA-β-gal) activity was more pronounced in a subset of RPE aneuploid cells (Fig 4C). To quantify senescence, we measured loss of the nuclear lamina protein lamin B1, a robust biomarker of senescence both *in vitro* and *in vivo* [24]. Aneuploid and control RPE cells or fibroblast lines were assessed by comparing lamin B1 levels between populations that either displayed or lacked micronuclei (MN+ or MN-; Fig 4D-F). In the vast majority of lines tested here, MN+ cells displayed a significant decrease in lamin B1 expression compared with MN-cells regardless of ploidy (Fig 4D-F). Of note, the only exception was primary cell line FIB^+21^ (Fig 4F) which may have reached replicative senescence *in vitro* despite our experimental procedures being performed at low passage. Together, these data suggest that the aberrant progeny of aneuploid cells undergo senescence irrespective of their ploidy status.

### Senescence bypass and accumulation of aberrant karyotypes after p53 depletion

Since p53 plays a major role in triggering senescence via multiple mechanisms including p21 induction [25], we next checked whether stabilization of p53 and p21 were observed in MN+ but not MN-populations of aneuploid and control cells (Fig 5A-C). Cells concomitantly stabilizing p53 and p21 were first identified by plotting the mean p53 intensity of each nuclear area against p21 level, and then classifying populations based on the presence or absence of a micronucleus (Fig 5B, C and Fig EV3; see cells in top right quadrant). With only two exceptions, all cell lines analyzed displayed significantly increased co-staining of p53 and p21 in MN+ cells relative to MN-cells (Fig 5B and C). Notably, western blot analysis revealed that RPE^+18+18^ displayed increased steady-state levels of p21 protein (Fig EV1G), likely accounting for the lack of differences observed when comparing p53/p21 levels between MN+ and MN-cells. Taken together, these results suggested that p53 may be activated in response to chromosome mis-segregation or DNA damage in the aberrant progeny of aneuploid cells, potentially leading to stabilization of p21 and senescence-associated growth restriction.

**Figure 5.**
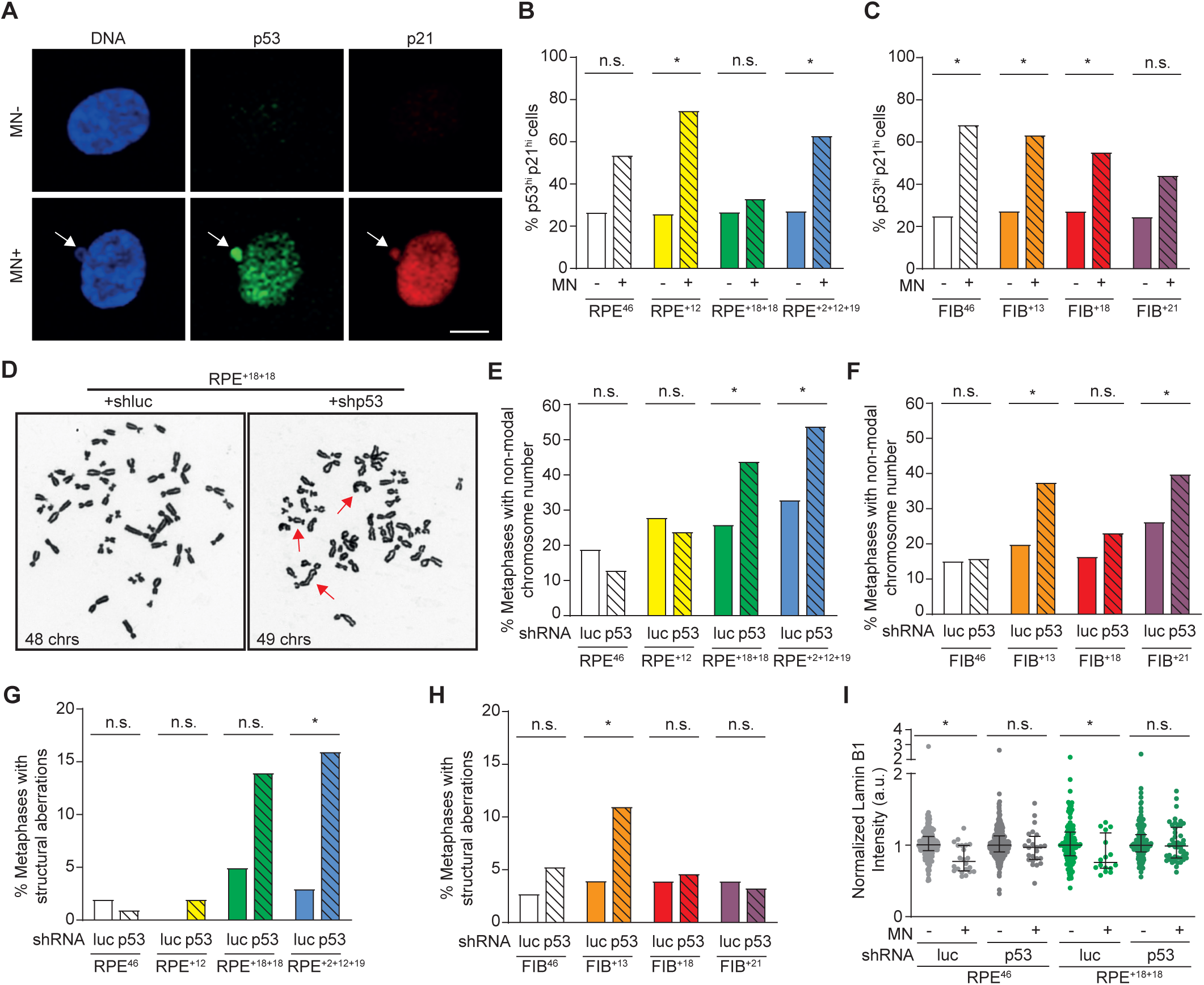
p53 restrains the growth of chromosomally unstable daughter of aneuploid cells. A Representative images of p53 and p21 immunofluorescent staining in RPE^+2+12+19^ cells with (MN+) and without (MN-) or micronuclei (white arrow). DNA was stained with DAPI (scale bar: 10µm). B,C Quantification of MN- and MN+ cells with high p53 and p21 staining in the indicated cell lines (*p<0.05 by Fisher’s exact test). Normalized p53 and p21 immunofluorescence levels per cell are reported in EV3. D Representative metaphase spreads of p53 knockdown (shp53) or control (shLuc) RPE^+18+18^ cells (red arrows: structural chromosome aberrations). E,F,G,H, Percentage of cells in the indicated lines carrying p53 knock down or control with (E-F) a chromosome number different from the modal karyotype (RPE^46^ and FIB^46^ = 46; RPE^+12^ and aneuploid fibroblasts = 47; RPE^+18+18^ = 48 and RPE^+2+12+19^ = 49) or with (G-H) structural chromosome aberrations (n=50 metaphase spreads from two independent experiments; *p<0.05 by Fisher’s exact test). I Normalized LaminB1 quantification in MN- and MN+ cells from p53 knockdown (shp53) or control (shLuc) RPE^WT^ or RPE^+18+18^ lines (n=16-46 MN+ cells; n=124-364 MN-cells). Error bars represent the median and interquartile ranges (*p<0.05 by Mann-Whitney test).

We next tested whether depletion of p53 in RPE and human fibroblast aneuploid clones would allow aberrant daughter cells to continue proliferation, thereby generating karyotypic variability in the total population (Fig 5D-H). Luciferase control (shluc) or p53 knockdown (shp53, Fig EV4) aneuploid and control cell lines were grown for two passages following viral transduction. Metaphase spreads were prepared at different time points to assess the extent of numerical and structural chromosome aberrations (Fig 5D-H). Down-regulation of p53 in diploid RPE^46^ and FIB^46^ cells was insufficient to generate karyotypic heterogeneity in short-term cultures, as evidenced by the lack of any significant change in chromosome copy numbers or structural features (Fig 5D-H) and consistent with the fact that these lines do not display any underlying chromosomal instability *in vitro* (Fig 2, 3 and 4A-B). In contrast, the chromosomally unstable aneuploid lines RPE^+18+18^, RPE^+2+12+19^, FIB^+13^ and FIB^+21^ (Fig 2, 3 and 4A-B), presented with a significant increase in metaphase spreads featuring either structural aberrations or chromosome numbers deviating from the mode (Fig 5D-H). Metaphase spreads prepared from RPE^+18+18^ cells are shown in Fig 5D to illustrate the concomitant change in chromosome number and structure, in line with reports that chromosome segregation errors generate DNA damage which can subsequently leading to structural aberrations [18]. These results suggest that the karyotypic heterogeneity observed after p53 knockdown is a consequence of the rescued viability of daughter cells arising from aberrant mitosis of aneuploid mother cells. In agreement with this concept, p53 depletion was sufficient to rescue the senescent phenotype of MN+ RPE^+18+18^ aneuploid cells (Fig 5I). Taken together, these results suggest a role for p53 in activating senescence following chromosome mis-segregation of aneuploid mother cells, thus preventing heterogeneity and genomic instability in the derivative population.

These data reveal that even when p53-dependent cell-autonomous mechanisms failed to restrain the growth of aneuploid cells, this pathway is still capable of inducing senescence and halting cell cycle progression of karyotypically unstable daughter cells. This finding suggests that both cell-autonomous and extrinsic mechanisms of growth inhibition should act in concert to ensure the effective clearance of unstable aneuploid cells, and are thus required in combination to protect multicellular eukaryotes against tumorigenesis.

## Materials and Methods

### Cell Culture

Cell lines 293T and hTERT-RPE1 (‘RPE’) were obtained from the American Type Culture Collection (Rockville, MD). Human primary aneuploid cell lines Fib^+13^ (GM00526), Fib^+18^ (GM03538), Fib^+21^ (GM04616) and diploid control (Fib^46^, GM01381) were purchased from the Coriell Repository. Cells were cultured in DMEM (293T), DMEM:F12 (RPE) or MEM (primary human fibroblasts) containing 10% FBS, 100 IU ml-1 penicillin and 100 μg ml-1 streptomycin (Gibco). All cell lines were maintained at 37 °C with 5% CO_2_ atmosphere, regularly tested for mycoplasma contamination, and used at low passage (≤10) to avoid selection in culture. In a subset of experiments, some cell lines were synchronised in S-phase with 2 mM thymidine (Sigma-Aldrich) for 24h, washed with PBS, then released in standard media or supplemented with the indicated compounds. To induce aneuploidization prior to single-cell separation, RPE cells were grown at high density for 72h to synchronize them in G1 by growth inhibition, then split into lower density for 20h after and subsequently exposed to 0.5 µM reversine for 6 h, unless otherwise stated. Following reversine treatment, cells were released into standard media and the following day these were single-cell flow-sorted into 96 well plates using a BD influx apparatus. Colonies grown from single cells were plated into 10cm dishes ∼2 weeks later and allowed to expand further. Karyotyping and cryopreservation was performed when the 10cm dishes reached confluence (∼20-23 cell doublings assuming no cell death).

### Retroviral and lentiviral infection

The retroviral pRetroSuper vector bearing the blasticidin resistance gene together with shRNA against luciferase (control) or p53 were kind gifts from Dr. Mathijs Voorhoeve. The p53 target sequence is described in [26]. Retroviruses were generated by transfecting 3×10^6^ 293T cells plated on 10cm dishes with 2.5 µg pCL ampho (retroviral packaging vector) and 7.5 µg pRetroSuper-shluc or -shp53 using Lipofectamine 2000 (Gibco) according to the manufacturer’s instructions. Virus-containing supernatants were harvested 48h after infection. A total of 2×10^5^ target cells were transferred into 6-well plates and infected with fresh or frozen retroviruses (diluted 1:1 with fresh medium) in the presence of 8 µg/ml polybrene. Cells were then selected with 5 µg/ml blasticidin for two weeks. H2B-mCherry and GFP-tubulin fusion open reading frames from pcDNA3-H2B-mCherry (a gift from Robert Benezra [Memorial Sloan Kettering Canecr Center, NY], Addgene plasmid # 20972) and pAcGFP1-Tubulin (Clontech) were subcloned into pLVX-Puro lentiviral vector (Clontech) using the BamH1 and XbaI restriction sites. Lentiviruses were generated by transfecting 293T cells with lentiviral vector (3.75 µg) and packaging plasmids (3.75 µg pMDLg/pRRE, 1.25 µg pRSV-Rev, 1.25 µg pCMV-VSV-G) using Lipofectamine 2000. Cell-free supernatants were harvested 48h after transfection and used for cell infection at 1:10 dilutions in the presence of 8 µg/ml polybrene. Cells were selected with 5 µg/ml puromycin for one week.

### RT-qPCR

Total RNA was extracted using the RNeasy mini kit (Qiagen) according to the manufacturer’s instructions. Reverse transcription of standardized RNA samples was performed using the SuperScript® III First-Strand Synthesis SuperMix kit (Life technologies) and mRNA copy numbers were determined by real-time quantitative PCR using the 7900HT Fast Real-Time PCR System (Applied Biosystems) or QuantStudio 7 Flex Real-Time PCR System (ThermoFisher Scientific). cDNA together with specific primers were mixed with PerfeCTa SYBR Green FastMix (Quantabio) following the supplier’s protocol. The oligonucleotides for each gene were as follows: p53: 5’-CAACAACACCAGCTCCTCTC and 5’-CCTCATTCAGCTCTCGGAAC, Actin: 5’-GGATCGGCGGCTCCAT and 5’-CATACTCCTGCTTGCTGATCCA.

### Western blotting

For western blotting, cells were re-suspended in ice-cold RIPA lysis buffer (ThermoFisher Scientific) supplemented with Complete Protease Inhibitors (Roche). Protein concentration in the lysate was determined using the Quick Start Bradford 1x Dye Reagent (Biorad), then 80-100 µg total protein was loaded into 4–15% Precast SDS-PAGE Gels (Biorad) and transferred onto nitrocellulose membranes (0.45 µm, Thermo Scientific). Blots were incubated in Odyssey blocking buffer (LI-COR) followed by addition of primary antibodies against p53 (DO-1, Abcam), p21 (12D1, Cell Signalling) and beta-actin (mAbcam 8226, Abcam). Fluorescently labeled secondary antibodies enabled subsequent protein measurement using the Odyssey Imaging system (LI-COR Biosciences).

### Quantification of cell death

After trypsinisation, the cell suspension was centrifuged at 1200rpm for 3mins, and then washed with PBS twice. The cell pellet was then incubated with FxCycle™ PI/RNase Staining Solution (cat #F10797, Invitrogen) for 15mins in the dark before FACS was performed using MACSQuant Analyzer Flow Cytometer (Miltenyi). FACS profiles were analyzed using FlowJo v10 software.

### Metaphase preparation, karyotyping and ploidy classification

Cells grown to ∼80% confluency were treated with 100 ng/ml Colcemid solution (Gibco) for 4-6h, collected by trypsinization and centrifuged at 1000 rpm for 10min. Cell pellets were re-suspended in 75mM potassium chloride solution and incubated for 15 min in a 37 °C waterbath. Next, a 1/10 volume of 3:1 methanol/acetic acid was added to the cells prior to centrifugation at 1000 rpm for 15 min. Cells were then fixed by resuspension in 3:1 methanol/acetic acid solution, incubated for 30min at room temperature, centrifuged at 1200 rpm for 5 min, and finally washed once more with fixative. Cells were re-suspended in a small volume of fixative, dropped onto clean glass slides and allowed to air dry. For chromosome counting, metaphase spreads were stained with Giemsa stain (Gibco) and acquired using the fully automated Metafer imaging platform (MetaSystems). Chromosome numbers (dicentric chromosomes were counted as two) and the presence of structural aberrations were scored manually using ImageJ. G-banded karyotype analysis was performed by the Cytogenetics Laboratory at the Genome Institute of Singapore. Spectral karyotyping was performed using SKY probes according to manufacturer’s instructions (Applied Spectral Imaging). Alternatively, multicolor FISH was performed using MetaSystem’s 24XCyte chromosome painting probes according to manufacturer’s instructions. To evaluate ploidy and chromosome instability a total of 20 metaphase spreads were used unless otherwise specified. A clone was classified diploid if modal chromosome number was 46 and aneuploid in all other cases. Polyploid cells with chromosome number >65 were seldom found.

### Fluorescence in situ hybridization (FISH)

To detect the presence of specific chromosomes, metaphase preparations (described above) or interphase cells were dropped onto glass slides and co-denatured with XA 13/18/21 AneuScore probe mix (MetaSystems) at 75 °C for 2 min before being placed in a humidified slide incubator (Eppendorf) at 37 °C for 24 h. Following hybridization, slides were washed first in 0.4X SSC (pH 7.0) at 72°C for 2 min, then in 2x SSC/0.05% Tween20 (pH 7.0) at room temperature for 30 sec, rinsed briefly in distilled water and mounted on microscope slides with fluorescence mounting media (Dako) together with DAPI. Cells were visualised using the automated Metafer imaging platform (MetaSystems). For FISH on binucleated cells, RPE and primary fibroblast cells were grown on coverslips and treated with 4uM dihydrocytochalasin B (Sigma-Aldrich) for 24-30 h before fixation. At the end of DCB treatment, cells were washed once with PBS and fixed with 3:1 methanol/acetic overnight at 4 °C. Coverslips were air-dried and hybridized with chromosome-specific probes as described above. Mis-segregation events were scored when an unbalanced chromosome copy number ratio was observed when comparing the two daughter nuclei. Only cases with an even number of total signals were included in the analysis. RPE+^2+12+19^ and control were synchronized with thymidine before DCB exposure.

### Immunofluorescence and Confocal microscopy

For immunostaining, cells grown on glass coverslips were fixed in 4% paraformaldehyde in PBS for 15 min at room temperature and permeabilized for 5 min in PBS containing 0.25% Triton-X 100. Cells were washed twice with PBS and then incubated in blocking solution (PBS with 5% normal goat serum, 2% BSA and 0.1% Triton-X 100) for 1h at room temperature. Cells were then incubated with primary antibodies diluted in blocking solution overnight at 4 °C, washed three times with PBS with 0.1% Triton-X 100 and incubated at room temperature for 1h with secondary antibodies diluted in blocking solution. After several wash steps, coverslips were briefly rinsed in distilled water and mounted on microscope slides in fluorescence mounting medium (Dako) with DAPI as nuclear counterstain. Primary antibodies used were as follows: rabbit anti-53BP1 (H-300, Santa Cruz), mouse anti-histone H2Ax (S139) (JBW301, Upstate), mouse anti-p53 (DO-1, Abcam) and rabbit anti-p21 Waf1/Cip1 (12D1, Cell Signaling). Alexa 488 conjugated goat anti-mouse and Alexa 568 goat anti-rabbit (Molecular Probes) secondary antibodies were used. Cells were visualised using an Olympus FV1000 inverted confocal microscope (Olympus Microscopes, Essex, UK) equipped with 405, 488 and 561 nm lasers for excitation and spectral/band-pass emission filters were used for acquisition of confocal images and construction of z-stacks. An Olympus Plan Apo × 40/1.0 oil immersion objective lens was used. Z-stacks with 0.5 µM steps were acquired and mean fluorescence intensity of lamin B1, p53 or p21 on DAPI-stained interphase nuclei was measured using a sum intensity projection in Fiji software (ImageJ, National Institutes of Health [NIH], Bethesda, MD) on all planes of the acquired stacks images. To allow for comparison among samples, the mean fluorescence intensity of each cell was normalized to the median fluorescence intensity value of each individual image.

### Time-lapse imaging

Cells expressing H2B-mCherry and GFP-tubulin were grown on 35 mm glass-bottom dishes (Ibidi) and maintained in an on-stage incubator (37 °C, 5% CO2 humidified atmosphere). Live cell imaging was performed using a spinning-disk confocal consisting of a motorized Nikon Ti inverted microscope (Nikon, Tokyo, Japan) equipped with a Plan Apo × 60/1.4 oil immersion objective lens, a 491 and 561 nm laser (FRAP-3D laser launch; Photometrics) and CSU-22 scanning head (Yokogawa Electric Corp., Tokyo, Japan), Evolve EM-CCD camera (Photometrics) and the Nikon Perfect Focus System. Imaging was controlled using MetaMorph software (50-100 ms exposure, EM-gain 300, 10% laser power, Molecular Devices, LLC, Sunnyvale, CA). Z-stacks (3 µm) covering the entire volume of the mitotic cells were collected every minute for 2-24 h depending on the experiment. All movies were analyzed using Fiji software (ImageJ, National Institutes of Health (NIH), Bethesda, MD).

### Statistical Analysis

Statistical significance was determined by Student’s t-test, Mann-Whitney test, two-tailed Fisher’s exact test as appropriate using GraphPad Prism 5 software (La Jolla, CA, USA). P<0.05 was considered statistically significant.

## Acknowledgements

We thank N. Pavelka for helpful discussions, M. Voorhoeve for constructs, S. Lim and S. Davila for G banding analysis (GIS, Singapore), J. Lim and G. Wright (IMB Microscopy Unit) for imaging analysis and GR lab members for technical help and suggestions. N. McCarthy of Insight Editing London reviewed the manuscript. The G.R. lab is supported by the NRF Investigatorship Award (NRF-NRFI05-2019-0008).

## Author contributions

M.G. and G.R. conceived and designed experiments; M.G., C.K.W. and G.R. prepared figures and wrote the manuscript; M.G., C.K.W., J.S.L. and M.S. performed experiments. O.D. contributed expertise to the senescence assays. G.R. supervised the study. All authors read the manuscript and agreed with its content.

## Supplementary Figure Legends

**Figure EV1.**
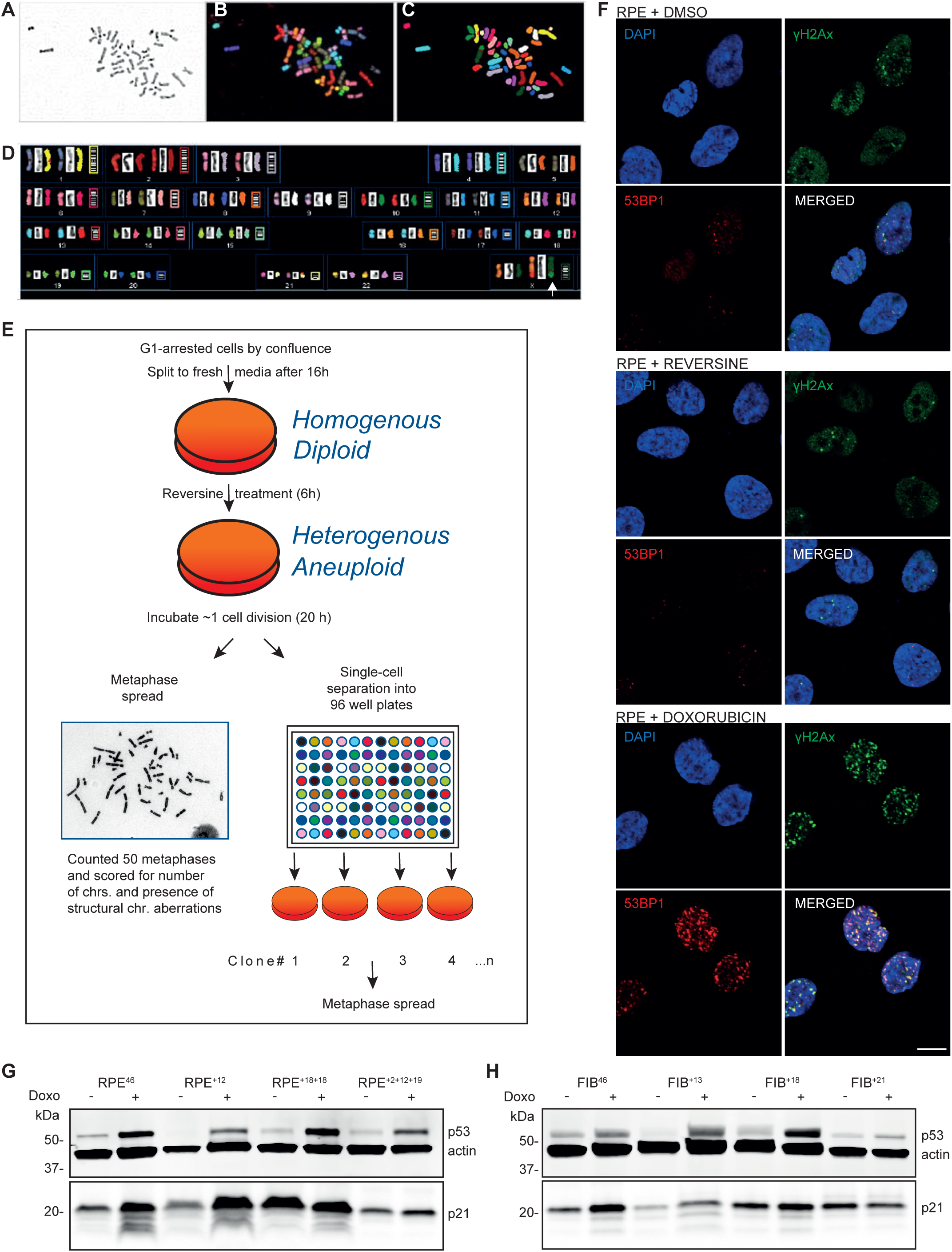
Spectral karyotype (SKY) analysis of RPEWT cells. A, B, C Inverted-DAPI (A), RGB (B) and classification pseudo-color (C) of a RPE cell treated with DMSO only. D Karyotype of the same metaphase spread. Arrow indicates a RPE characteristic X chromosome derivative, add(X)(q28). E RPE cell line was treated either with the MPS1 kinase inhibitor reversine or vehicle only then released into standard culture media for 20 h to allow acquisition of aneuploid cells (see Material and Methods for details). For each culture, a sample of the total cell population was used to determine overall aneuploidy induction (Fig 1A and B) while the remaining cells were single-cell sorted into 96-well plates to allow clonal amplification. After ∼20 generations (∼2-3 weeks), single-cell clones were expanded and their chromosome content was analyzed (Figure 1 D-G). F Immunolabeling with γH2Ax (green), 53BP1 (red) and DNA (DAPI, blue) in RPE cells after treatment with DMSO vehicle control (top panel), reversine (0.5µM, 6h, middle panel) or doxorubicin (300nM, 6h, bottom panel). Scale bar: 10µm. G, H Western blots of p53 and p21 in doxorubicin-treated (400nM 4h, +) or untreated (-) euploid or aneuploid RPE (G) or fibroblast (H) cell lines (ß-actin: loading control). Labels on the left indicate molecular weight markers in kilodaltons (kDa).

**Figure EV2.**
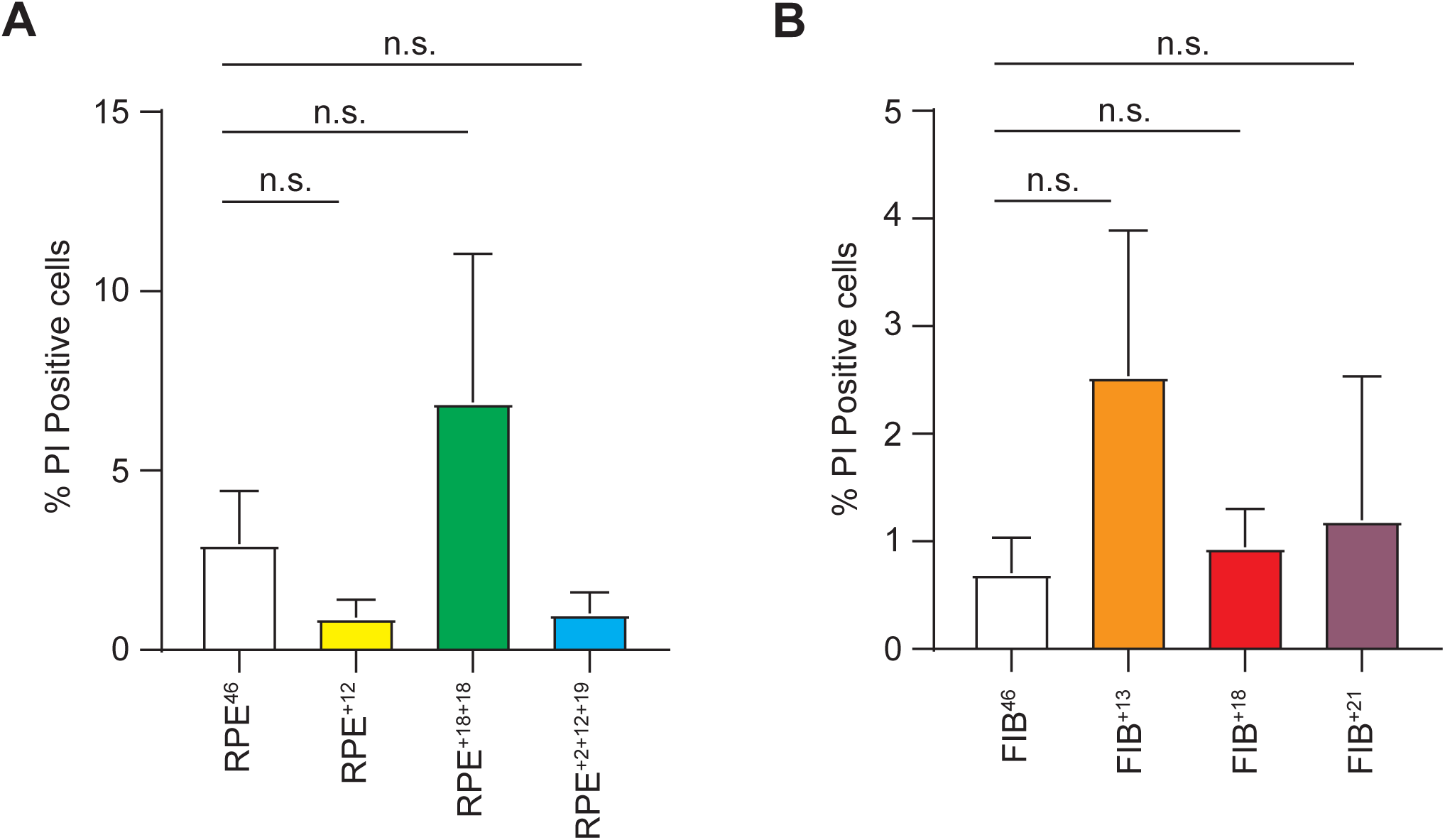
Aneuploid lines do not show increased cell death. A, B Quantification of apoptotic cells in euploid and aneuploid RPE (A) or fibroblast (B) cell lines. Percentage of Propidium Iodide positive cells was quantified by FACS. Three independent biological replicates were quantified and and sd are shown in the graphs. At least 60000 events were acquired for each biological replicate.

**Figure EV3.**
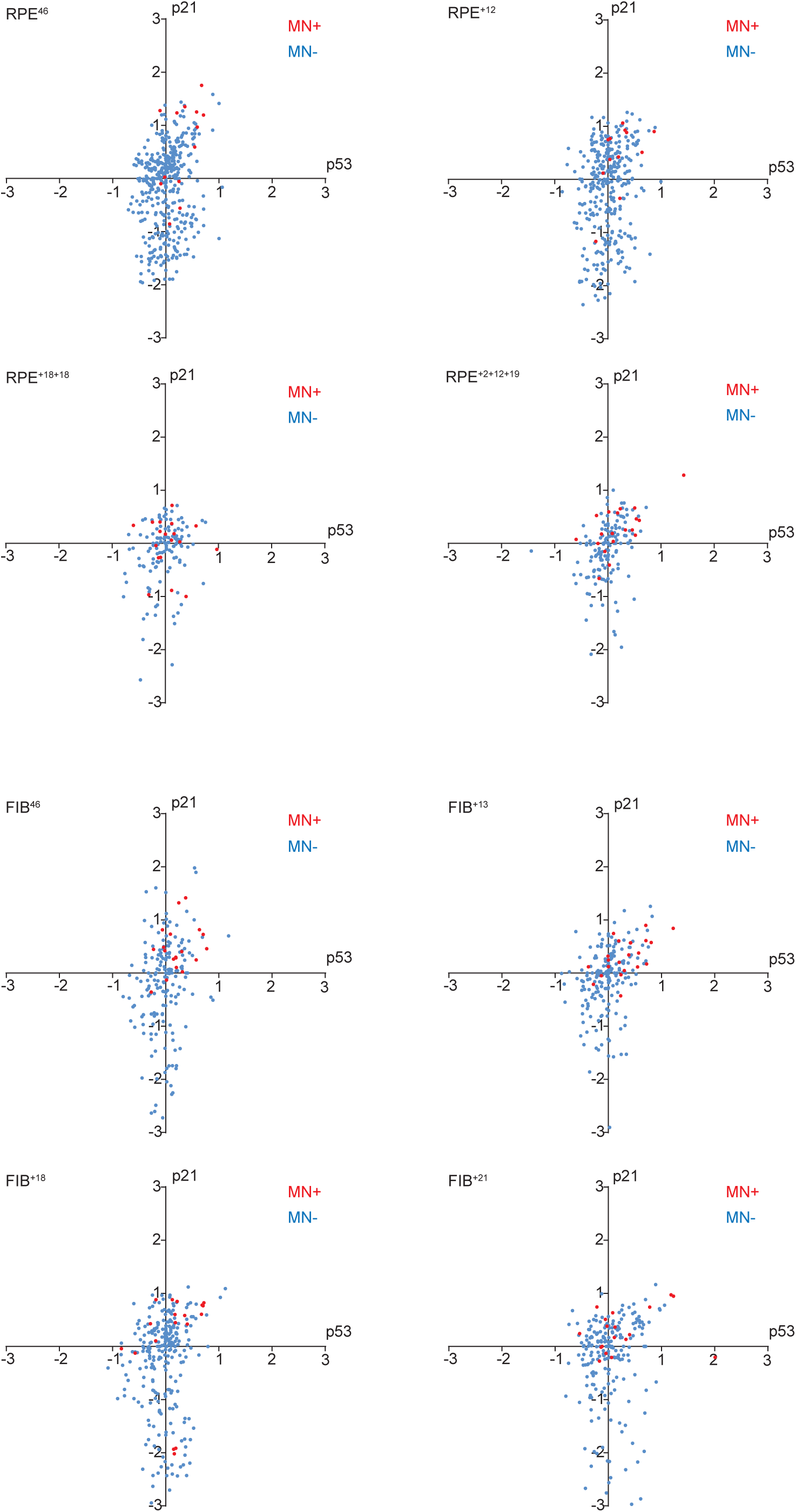
Quantification of p53 and p21 abundance in MN-versus MN+ cells. Plots of log_10_[normalised p53 intensity] (x-axis) against log_10_[normalised p21 intensity] (y-axis) in the indicated cell lines. Each dot represents an individual cell and MN- and MN+ populations are represented as blue and red dots, respectively (n=12-22 MN+ cells and n=144-411 MN-cells per cell line). For each line, the proportion of MN+ and MN-cells displaying elevated p53 and p21 expression is derived from the top-right quadrant and has been plotted in Fig 5B and C.

**Figure EV4.**
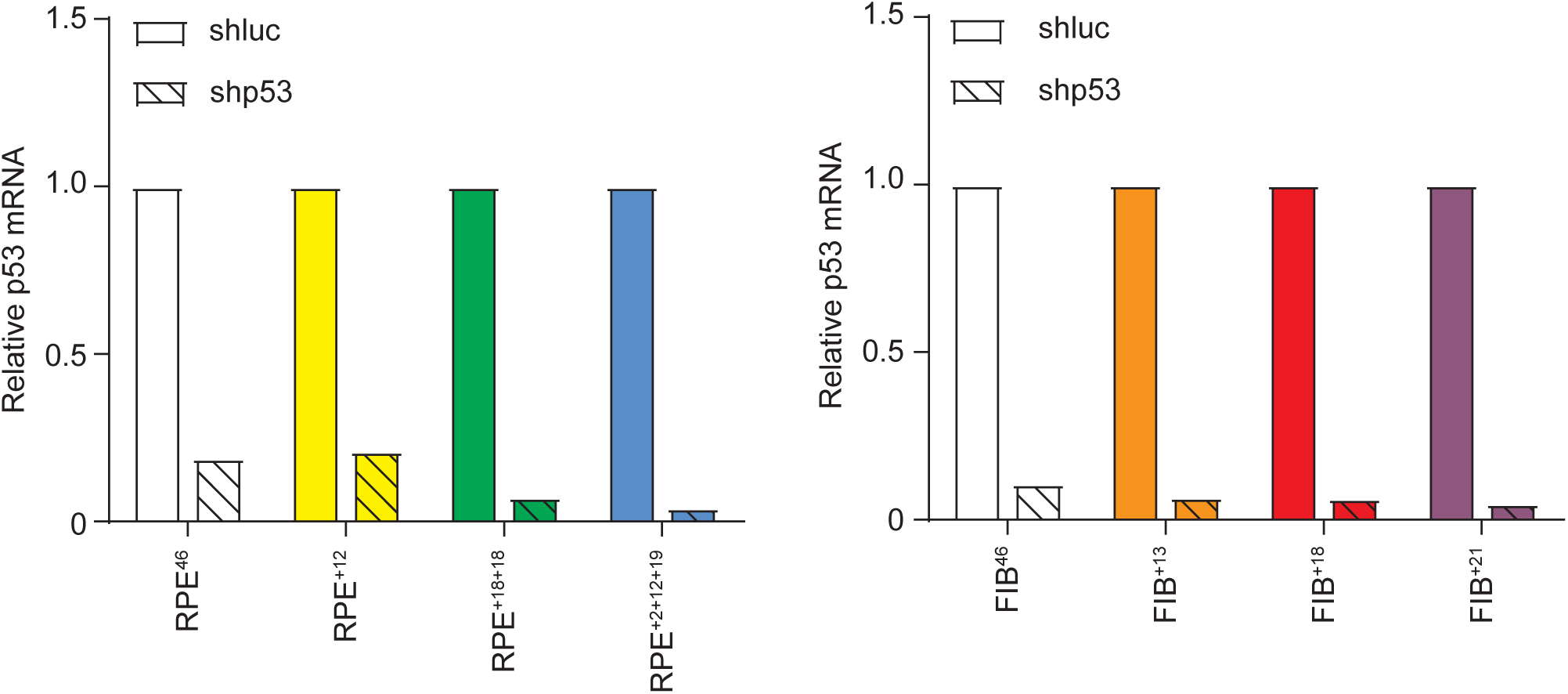
Validation of p53 knockdown in indicated cell lines. mRNA expression levels were quantified using qRT-PCR in the indicated cell lines. Solid: control (shLuc); Pattern p53 knockdown (shp53). ß-Actin was used for the normalization of expression levels.

## Supplementary Video Legends

**Movie EV1. Chromosome segregation of RPE+18+18 cell line.** Time-lapse imaging of a RPE^+18+18^ cell stably expressing H2B-mCherry. (A) equal chromosome segregation and (B) aberrant chromosome segregation (refer also to Fig 2A). Cells were recorded at 63X magnification every minute. Time=h:min. Movie is played at 6 frames per second.

